# Metabolic energy expenditure during level, uphill, and downhill running

**DOI:** 10.1101/2025.06.05.658094

**Authors:** David P. Looney, Wouter Hoogkamer, Rodger Kram

**Author notes:** Corresponding Author David P. Looney.

## Abstract

**Purpose:** To extend the Running Energy Expenditure Estimation (RE3) model for predicting metabolic rates during uphill and downhill running as well as to enhance the Hoogkamer-Taboga-Kram (HTK) equation for estimating metabolic rates during level and uphill running.

**Methods:** We combined running metabolic data from an original dataset (n=63) with individual subject data from 26 studies (n=424) and group mean data from 12 studies (n=187). Using this integrated dataset, we derived a new graded running term for the empirical RE3 model and updated the HTK equation coefficients for level and uphill running. We then compared the accuracy and precision of these new equations with the established American College of Sports Medicine (ACSM) and Minetti et al. equations based on the root-mean-square deviation (RMSD).

**Results:** Accuracy and precision of estimating level Ṁ were high for the Minetti et al. (RMSD, 1.44 W·kg^-1^), HTK (1.30 W·kg^-1^), and RE3 (1.27 W·kg^-1^) equations but much worse for the ACSM equation (1.82 W·kg^-1^). Agreement on uphill slopes was highest for the HTK (RMSD, 1.45 W·kg^-1^) and RE3 (1.41 W·kg^-1^) equations with less precision noted for the ACSM (2.17 W·kg^-1^) and Minetti et al. (2.18 W·kg^-1^) equations. When estimating Ṁ during downhill running, the RE3 equation performed marginally better (RMSD, 1.45 W·kg^-1^) than the Minetti et al. equation (1.57 W·kg^-1^).

**Conclusion:** The improved RE3 and HTK equations estimate metabolic rates during level and graded running with improved accuracy and precision. We provide a publicly available web-based metabolic rate calculator that simplifies estimation for researchers, practitioners, and runners alike.

## Introduction

Many people who run for health and fitness strive to *maximize* their rates of “burning calories” as an aid to weight management (Caudwell et al., 2014). In contrast, competitive runners seek to *minimize* their rate of metabolic energy expenditure (Ṁ) so that they can run faster at their physiological limits (Kipp et al., 2019). Military personnel and ultramarathon runners need to estimate their Ṁ so that they can consume adequate amounts of food for optimal performance during missions or races respectively. Many equations exist that can predict Ṁ during level running (Hall et al., 2004; Looney et al., 2023; Rubenson et al., 2007) and thus provide information relevant to these needs. Notably, we and our colleagues recently developed a generalized equation for estimating Ṁ during level running for the Running Energy Expenditure Estimation (RE3) model based on data compiled from over 50 studies (Looney et al., 2023). However, many runners also traverse a variety of uphill and downhill terrain. As any runner knows, uphill running is more strenuous, and downhill running is less so. But there are fewer equations that predict how much greater Ṁ is during uphill running and even fewer equations for how much Ṁ changes during downhill running.

Since 1975, the American College of Sports Medicine (ACSM) has espoused a simple equation for predicting oxygen uptake (V̇O_2_) during level and uphill running (Ozemek et al., 2025). The ACSM equation assumes that V̇O_2_ during uphill running is equal to the sum of V̇O_2_ for resting metabolism (3.5 mL·kg^-1^·min^-1^) plus the V̇O_2_ for running on the level and the V̇O_2_ required to lift the body mass vertically against gravity. The third term in the equation (vertical lifting) essentially assumes an unphysiologically high efficiency value (> 50%) (see Appendix). Given that isolated muscle efficiency (Barclay & Curtin, 2023) and muscular efficiency during cycling exercise (Ettema & Lorås, 2009) are both about 25%, using a value > 50% is in conflict with established physiology.

The pioneering efforts of Minetti et al. (2002) introduced a 5^th^ order polynomial equation based on physiological measurements that predicts the amount of metabolic energy expended during running per unit distance (J·kg^-1^·m^-1^, aka metabolic cost of transport, CoT) on uphill, level, and downhill slopes (referred to as the Minetti equation henceforth). However, the Minetti equation neglects the costs of generating force to support body weight, brake and propel the center of mass (COM) (Arellano & Kram, 2014). Further, the predictions of the Minetti equation diverge substantially from metabolic data measured during very steep uphill running (Giovanelli et al., 2016).

Hoogkamer et al. (2014) addressed the ACSM efficiency issue by accounting for differences in the biomechanics between uphill and level running. The Hoogkamer et al. (2014) approach (which we refer to as HTK, based on the last name initials of the authors) breaks the metabolic cost of uphill running into several constituents: the cost of bouncing perpendicular to the running surface (including the cost of swinging the arms and legs), the cost of running parallel to the surface, and the cost of producing mechanical work to lift the center of mass against gravity. At a given running speed on the level, Ṁ is directly proportional to the average ground reaction force (GRF) perpendicular to the surface and inversely proportional to the time of foot-ground contact (t_c_) (Kipp et al., 2018b; Kram & Taylor, 1990; Roberts et al., 1998). Since Hoogkamer et al. (2014) found that neither the perpendicular force nor t_c_ changed appreciably on moderate inclines, and since stride frequency changes little during uphill running, they assumed the combined cost of perpendicular bouncing plus limb swing remains nearly the same.

However, the GRF parallel to the running surface changes dramatically during uphill running. The braking impulse diminishes to near zero when running up a 9° incline at which slope the propulsive impulse only acts to advance the center of mass up the slope. Thus, Hoogkamer et al. (2014) modelled the cost of parallel running as decreasing exponentially with incline. Having accounted for the costs of generating perpendicular and parallel forces, Hoogkamer et al. (2014) could then properly consider the metabolic cost of the mechanical work required to lift the center of mass vertically. Fitting their equation to the metabolic data revealed that a constant delta efficiency for doing vertical work (29%) was then an appropriate assumption. We note that the HTK model was only developed and tested on moderate inclines (< 9°) and slow speeds (< 3 m·s^-1^) and for a modest number of participants (n = 8). Further, the HTK model predicted the metabolic cost of transport (J·kg^-1^·m^-1^) whereas most runners are concerned with Ṁ.

Recently, U.S. Army researchers derived a new empirical method to precisely estimate Ṁ during level, uphill, and downhill *walking* using a unique formula (Looney et al., 2019b). A graded walking factor was incorporated into an existing level walking equation (Looney et al., 2019a) and outperformed existing alternatives during cross-validation (Looney et al., 2019b). Given the similar “J-shape” relationships between grade and metabolic costs of transport for both walking and running (Minetti et al., 2002), deriving an empirical, graded running factor with a similar formula for the RE3 model is a logical next step.

Our primary goal was to extend the RE3 model as per Looney et al. (2019b) to predict Ṁ during level, uphill and downhill running across a range of speeds using a large database aggregated from multiple sources. A secondary goal was to first transform the HTK equation from CoT units to Ṁ units and then update the coefficients, using the same large database, to provide a generalized equation consistent with our present understanding of level and uphill running biomechanics and muscle physiology. Unfortunately, muscle function during downhill running remains poorly understood, and thus we presently cannot confidently develop a predictive equation for downhill running based on physiology. For ease of use, we aimed to provide readers with web-based Ṁ calculator that included the newly derived methods to support practical applications (University of Massachusetts Integrative Locomotion Lab., 2025).

## Methods

### Experimental design

Like our previous study (Looney et al., 2023), we combined running metabolic data from three types of sources: original individual subject data, published individual subject data (Table 1), and published group mean data (Table 2). We aggregated data from level, uphill, and downhill running trials to derive the new graded running term of the empirical RE3 model. Data from the level and uphill running trials was subsequently used to update the coefficients of the HTK equation. The accuracy and precision of Ṁ estimates were subsequently compared between these equations along with the existing equations from the ACSM (level and uphill running only) and Minetti et al. (2002).

**Table 1.** Participant characteristics (mean ± SD) plus treadmill grade and speed (min, max) for published individual participant datasets. f, number of female participants, n, number of participants.

| Reference | n (f) | Age (y) | Height (m) | Mass (kg) | Grade (%) | Speed (m·s<sup>-1</sup>) |
| --- | --- | --- | --- | --- | --- | --- |
| Balducci et al. (2017) | 26 (0) | 42 ± 10 | 1.77 ± 0.04 | 72 ± 6 | [0.0, 10.0] | [1.66, 3.33] |
| Balducci et al. (2023) | 15 (0) | 30 ± 9 | 1.78 ± 0.06 | 65 ± 8 | [0.0, 15.0] | [1.67, 3.33] |
| Bertschy et al. (2022) | 14 (5) | 23 ± 5 | 1.72 ± 0.14 | 68 ± 10 | [−5.0, 0.0] | 3.89 |
| Besson et al. (2023) | 41 (20) | 33 ± 7 | 1.70 ± 0.10 | 60 ± 10 | [0.0, 10.0] | [2.78, 3.89] |
| Breiner et al. (2019) | 19 (0) | 29 ± 6 | 1.77 ± 0.07 | 68 ± 8 | [−5.0, 7.5] | [2.78, 4.85] |
| Brill and Kram (2021) | 10 (0) | 29 ± 6 | 1.79 ± 0.06 | 68 ± 5 | [0.0, 26.8] | [1.20, 2.30] |
| Davies et al. (1974) | 1 (0) | 31 | 1.78 | 61 | [−45.0, 5.0] | [1.58, 4.68] |
| Ehrström et al. (2018) | 9 (0) | 39 ± 8 | 1.73 ± 0.06 | 68 ± 6 | [0.0, 10.0] | [2.50, 3.88] |
| Hoff et al. (2016) | 7 (0) | 24 ± 6 | 1.81 ± 0.06 | 72 ± 6 | 1.5 | [2.78, 4.92] |
| Hoogkamer et al. (2014) | 8 (4) | 26 ± 4 | 1.74 ± 0.12 | 67 ± 12 | [1.7, 14.1] | [2.00, 3.00] |
| Lemire et al. (2022) | 8 (0) | 29 ± 10 | 1.74 ± 0.03 | 62 ± 5 | [−15.0, 15.0] | [1.42, 5.83] |
| Lemire et al. (2023) | 19 (6) | 34 ± 10 | 1.75 ± 0.10 | 69 ± 12 | [−10.0, 10.0] | 2.78 |
| Lussiana et al. (2013) | 14 (0) | 23 ± 4 | 1.78 ± 0.05 | 70 ± 5 | [−8.0, 8.0] | 2.78 |
| Margaria (1938) | 4 (0) | 26 ± 6 | 1.78 ± 0.07 | 72 ± 4 | [−35.0, 15.0] | [0.73, 4.28] |
| Ogueta-Alday et al. (2018) | 48 (0) | 32 ± 7 | 1.76 ± 0.05 | 70 ± 7 | 1.0 | [2.25, 5.78] |
| Perrin et al. (2023) | 20 (0) | 28 ± 8 | 1.75 ± 0.05 | 65 ± 4 | [0.0, 15.0] | [2.22, 4.44] |
| Sabater Pastor et al. (2021) | 46 (17) | 36 ± 8 | 1.74 ± 0.09 | 65 ± 11 | [0.0, 15.0] | [1.33, 3.53] |
| Sabater Pastor et al. (2023) | 17 (0) | 29 ± 7 | 1.77 ± 0.06 | 64 ± 5 | [0.0, 10.0] | [2.78, 3.89] |
| Seki et al. (2020) | 12 (0) | 22 ± 1 | 1.71 ± 0.05 | 60 ± 4 | [−6.0, 6.0] | 3.61 |
| Sundström et al. (2021) | 6 (0) | — | 1.78 ± 0.06 | 71 ± 8 | [−17.6, 0.0] | 3.33 |
| Vercruyssen et al. (2016) | 13 (0) | 38 ± 5 | 1.76 ± 0.05 | 68 ± 6 | [0.0, 10.0] | [2.50, 2.77] |
| Vercruyssen et al. (2017) | 12 (0) | 40 ± 5 | 1.75 ± 0.05 | 68 ± 6 | [0.0, 10.0] | [2.50, 3.89] |
| Vernillo et al. (2015) | 14 (0) | 40 ± 7 | 1.77 ± 0.07 | 73 ± 7 | [−5.0, 5.0] | 2.78 |
| Vernillo et al. (2020) | 14 (0) | — | — | — | [−5.0, 5.0] | 2.80 |
| Whiting et al. (2022) | 16 (0) | 27 ± 5 | 1.79 ± 0.05 | 67 ± 4 | [−5.2, 5.2] | 3.61 |
| Willis et al. (2019) | 11 (6) | $34 \pm 4$ | $1.70 \pm 0.05$ | $62 \pm 7$ | [0.0, 12.0] | [2.50, 4.17] |

**Table 2.** Participant characteristics (mean ± SD) plus treadmill grade and speed (min, max) for published group mean datasets. f, number of female participants, n, number of participants.

| Reference | n (f) | Age<br>(y) | Height<br>(m) | Mass<br>(kg) | Grade<br>(%) | Speed<br>(m·s<sup>-1</sup>) |
| --- | --- | --- | --- | --- | --- | --- |
| Balsalobre-Fernández et al. (2018) | 12 (0) | 26 ± 5 | 1.79 ± 0.05 | 67 ± 8 | 1.0 | [4.17, 5.56] |
| Barnes and Janecke (2017) | 15 (8) | 22 ± 2 | 1.73 ± 0.08 | 61 ± 8 | 1.0 | [3.58, 4.92] |
| Filardo et al. (2008) | 36 (0) | 24 ± 3 | 1.78 ± 0.06 | 73 ± 8 | [0.0, 5.0] | [2.00, 2.67] |
| Gimenez et al. (2014) | 11 (0) | 37 ± 8 | 1.78 ± 0.06 | 69 ± 7 | [-8.0, 8.0] | 2.50 |
| Giovanelli et al. (2016) | 15 (10) | 33 ± 8 | 1.75 ± 0.09 | 64 ± 9 | [16.6, 81.6] | [0.55, 2.14] |
| Helgerud et al. (1990) | 12 (6) | 26 ± 3 | 1.74 ± 0.08 | 67 ± 10 | 1.7 | [2.33, 4.33] |
| Jones and Doust (1996) | 9 (0) | 25 ± 5 | — | 72 ± 3 | [0.0, 3.0] | [2.92, 5.00] |
| Lemire et al. (2021) | 29 (10) | 34 ± 10 | 1.74 ± 0.09 | 68 ± 12 | [-20.0, 20.0] | [2.22, 3.89] |
| Ortiz et al. (2017) | 11 (4) | 31 ± 8 | 1.71 ± 0.08 | 67 ± 9 | 57.7 | [0.30, 0.90] |
| Paes and Fernandez (2016) | 10 (10) | 29 ± 8 | 1.78 ± 0.07 | 78 ± 6 | 1.0 | [2.46, 3.16] |
| Snyder and Farley (2011) | 9 (0) | 29 ± 4 | — | 69 ± 5 | [-5.2, 5.2] | 2.80 |
| Sundby and Gorelick (2014) | 18 (18) | 28 ± 5 | 1.68 ± 0.08 | 61 ± 8 | 1.0 | [2.68, 3.35] |

### Original individual subject data

Original individual subject data were collected from 62 United States Army civilians and Soldiers (10 women, 52 men; age, 24 ± 7 y; height, 1.74 ± 0.08 m; body mass, 77 ± 15 kg). Level running data were previously analyzed from 45 participants from this group (Looney et al., 2023). After completing a ∼ 20 min warm-up walk, participants ran for 3-4 min on a motorized treadmill while wearing standard physical training attire (i.e., shorts, t-shirt, socks, running shoes) on two separate visits. The treadmill was initially set to approximately 70.0-77.5% of the individual’s self-reported two-mile (3.2 km) run pace at 0% incline for the first stage with incremental increases in grade (2.5%) or speed (7.5% two-mile run pace) every 2 min stage afterward (Figueiredo et al., 2022; Schafer et al., 2025). Oxygen uptake (V̇O_2_) and carbon dioxide production (V̇CO_2_) were measured over the final minute of the run with a metabolic measurement system (TrueOne 2400; ParvoMedics, Sandy, UT, USA). The study was approved by the governing Institutional Review Board. Participants gave their voluntary informed consent after a briefing on the purpose of the study and potential risks. Investigators adhered to Department of Defense Instruction 3216.02 and Title 32 Code of Federal Regulations 219 on the use of volunteers in research. The study was performed in line with the principles of the Declaration of Helsinki.

### Published individual participant data

We searched PubMed, Web of Science, and Google Scholar for published individual participant datasets from studies that measured steady-state treadmill running metabolic energetics in apparently healthy men and women (18–44 y) via indirect calorimetry. Search terms included “treadmill”, “running”, “grade OR uphill OR downhill”, “energy OR metabolic OR oxygen”, and “cost OR economy OR expenditure OR rate.” We reviewed the retrieved publications reference lists and contacted key authors in the field to find additional studies. Authors of the current study also contributed individual data they collected or had been shared with them previously. We excluded data from individuals who reported conditions that would contraindicate intense running exercise (cardiovascular defects, chronic obstructive pulmonary disease, diabetes mellitus, musculoskeletal defects or disorders, thyroid disorders, etc.). Additionally, trials were excluded if they involved external loading (e.g., backpacks, weighted vests), unusual equipment (e.g., biohazard protection suit, exoskeleton), environmental stressors with pronounced metabolic effects (e.g., cold, heat), reported respiratory exchange ratio (RER) values less than 0.7 or greater than 1.0, or reported blood lactate concentrations > 4 mmol·L^-1^. Table 1 displays characteristics of the 424 participants (58 women) from the 26 individual participant data studies.

### Published group mean data

Group mean datasets were aggregated using essentially the same procedures as outlined for the individual participant datasets. As additional inclusion criteria, the sample size (n), mean, and standard deviation (SD) were required for group mean dataset trials to be included. Table 2 shows characteristics of the 187 participants (66 women) from the 12 group mean data studies included in our analysis.

### Running Energy Expenditure Estimation (RE3) Model

The level running equation for the RE3 model can be written as:

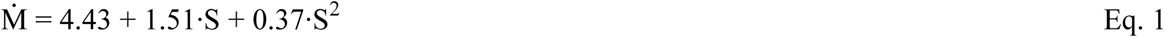

where Ṁ is rate of metabolic energy expenditure (W·kg^-1^) and S is running speed (m·s^-1^). We sought to add an empirical graded term to Eq. 1 in order to model the “J-shaped” relationship between surface grade and Ṁ (Looney et al., 2019b). Critical characteristics of this “J-shaped” relationship include the increase Ṁ on positive slopes (Dill, 1965) that remains linear even at extreme uphill grades (Giovanelli et al., 2016); the decrease in Ṁ on gradual downhill grades as propulsive force requirements are supplied by gravity (Gottschall & Kram, 2005; Margaria et al., 1963); the minimization of running Ṁ between −10 to −20% grade (Lemire et al., 2023; Margaria, 1938; Minetti et al., 2002); and the eventual upturn in Ṁ at steeper negative grades presumably due to the additional eccentric braking forces required to maintain a constant running speed against the greater parallel component of gravitational force (Davies et al., 1974; Margaria, 1938; Minetti et al., 2002). The RE3 Graded Running Equation for estimating Ṁ from running speed (S, m·s^-1^) and grade (G, decimal rise/run) can be written as:

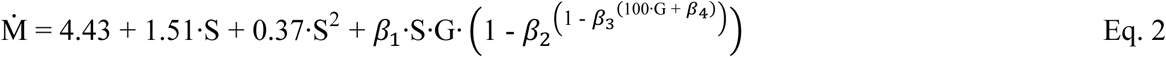

where β_1_, sets the maximum rate of Ṁ increase for positive grades; β_2_, sets the maximum rate of Ṁ increase for negative grades; β_3_, sets the grade and value of the minimal Ṁ; and β_4_, sets the negative grade at which Ṁ returns to level running values.

### Hoogkamer-Taboga-Kram (HTK) Model

The original HTK equation (Hoogkamer et al., 2014) predicts the net metabolic cost of transport (Net CoT) and consists of three terms:

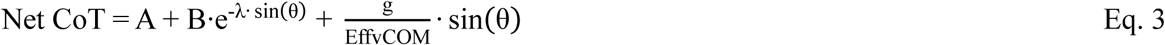

where the first term represents the metabolic cost of bouncing perpendicular to the running surface (including the cost of swinging the arms and legs), the second term represents the cost of running parallel to the surface, which decays exponentially with decay constant λ as a function of the sine of the incline (θ) in degrees, and third term represents the cost of producing mechanical work to lift the center of mass against gravity (g) with efficiency Eff_vCOM_. Here, “net” refers to the subtraction of the standing Ṁ from the gross Ṁ prior to calculating CoT. In the original HTK equation, the decay constant of the cost of running parallel to the surface is based on the decay constant of the experimentally measured braking impulses and independent of the metabolic data the equation was fitted to. Therefore, we adopted that value of 18.24 here. We then multiplied the HTK equation for Net CoT by running speed (S, m·s^-1^) to obtain net Ṁ, and added the generalized standing Ṁ determined by Looney et al. (2019a) (1.44 W·kg^-1^) to obtain the gross Ṁ. Coefficients A, B and Eff_vCOM_ were recalculated by fitting this equation to the large compiled database of metabolic data.

### Statistical analysis

We used R Version 4.12; R Foundation for Statistical Computing; Vienna, Austria) (R Core Team, 2014) to analyze all data and create each figure. Data are reported as mean ± SD unless stated otherwise. Rates of metabolic energy expenditure were determined from measured V̇CO_2_ and V̇O_2_ with an indirect calorimetry equation (Kipp et al., 2018a) derived from a previously developed nonprotein respiratory quotient table (Péronnet & Massicotte, 1991). Missing individual RER values were assumed to be equal to the trial mean if reported or 0.90 when no value was reported. Outliers were screened quantitatively using the 3 median absolute deviations from the median approach (Leys et al., 2013).

Equation coefficients were determined using nonlinear mixed effects modeling with the “nlme” package (Pinheiro et al., 2017) with random effects of group within study to account for the dependence among repeated measures. Datapoints were weighted in the regression by n size, with each individual participant coded as n = 1. We used linear mixed effects models, derived using the “lme4” package (Bates et al., 2014), to assess as accuracy and precision of Ṁ estimates using similar statistical procedures as our previous study (Looney et al., 2023). Accuracy was assessed as the bias weighted by n size and calculated as the mean paired difference between estimated and measured Ṁ. Precision was assessed as the SD of paired differences weighted by n size and calculated as the root-sum of the mean squared difference between estimated Ṁ and the overall bias plus the within-group variance. The root-mean-square deviation (RMSD), as an indicator of combined accuracy and precision as well as overall agreement, was calculated as the square root of the sum of the squared bias and SD of paired differences.

## Results

Fig1 shows the J-shaped relationship between grades within ± 0.45 (rise/run, i.e., 45%) and the net CoT using the compiled dataset of individual and group mean data. The RE3 Graded Running Equation was determined to be:

**Fig 1.**
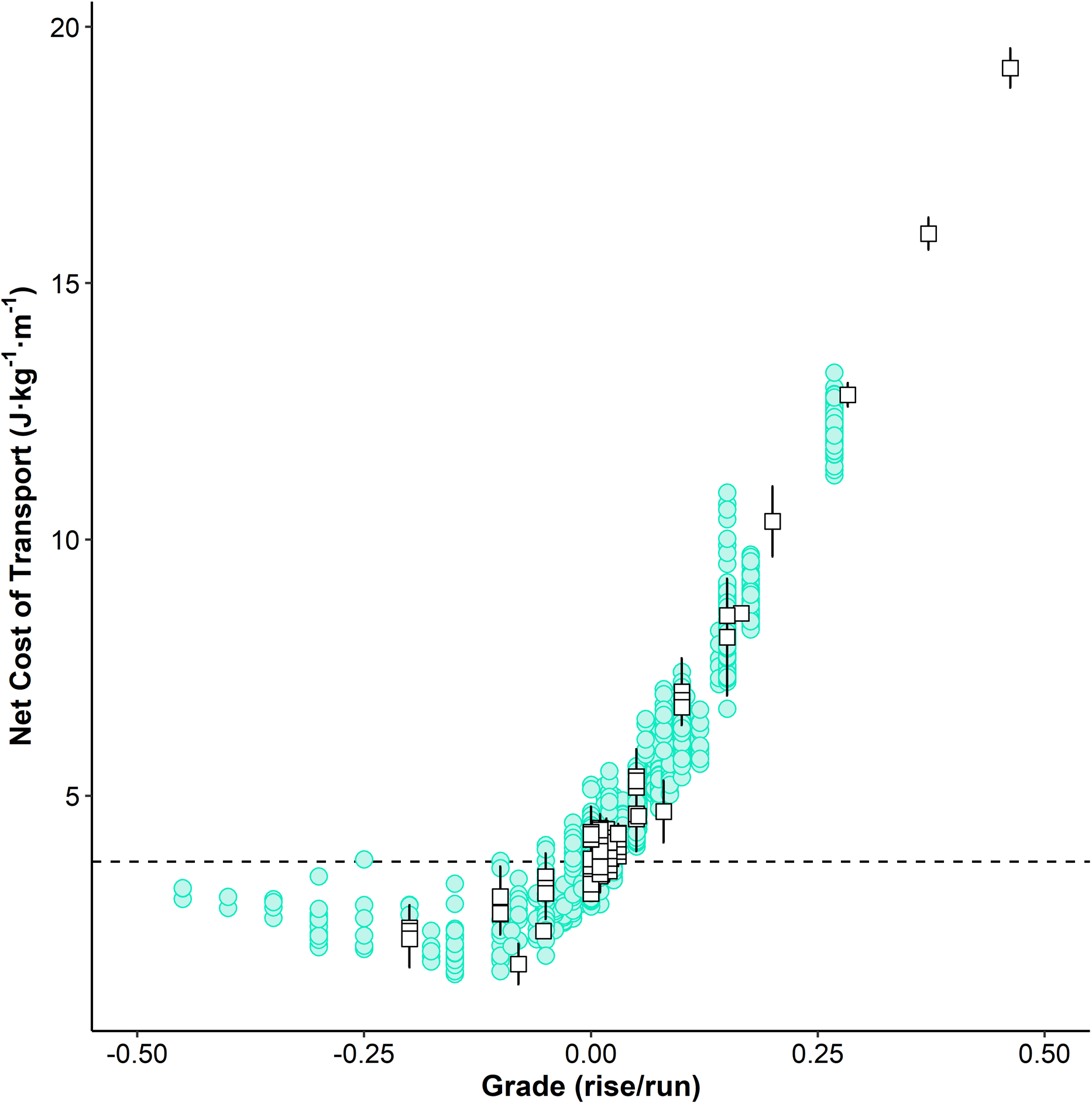
Net cost of transport, calculated as the difference between the measured metabolic rate and an assumed constant standing metabolic rate (1.44 W·kg^-1^) (Looney et al., 2019a) divided by speed, during running on decimal grades between −0.45 and 0.45 for individual subject data (green circles) and group mean data (white squares) ± standard deviation. Dashed line, mean net cost of transport when grade = 0 (3.71 W·kg^-1^).

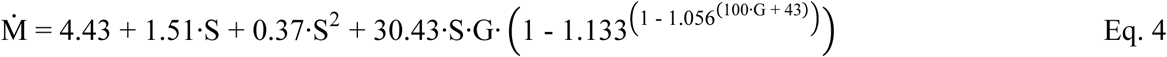

Fitting the HTK model coefficients to the aggregated dataset resulted in the following equation:

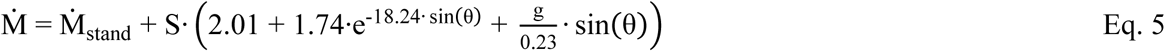

Table 3 displays the accuracy and precision of each of the equations when estimating Ṁ during level, uphill, and downhill running. Relative accuracy and precision of estimating level Ṁ were high for the Minetti, HTK, and RE3 equations while the ACSM equation tended to substantially overestimate Ṁ (Fig2 and Table 3). The HTK and RE3 equations had the highest agreement on uphill slopes with less precision noted for the ACSM and Minetti equations (Fig3). The RE3 equation performed marginally better than the Minetti equation when estimating Ṁ during downhill running (Fig4).

**Fig 2.**
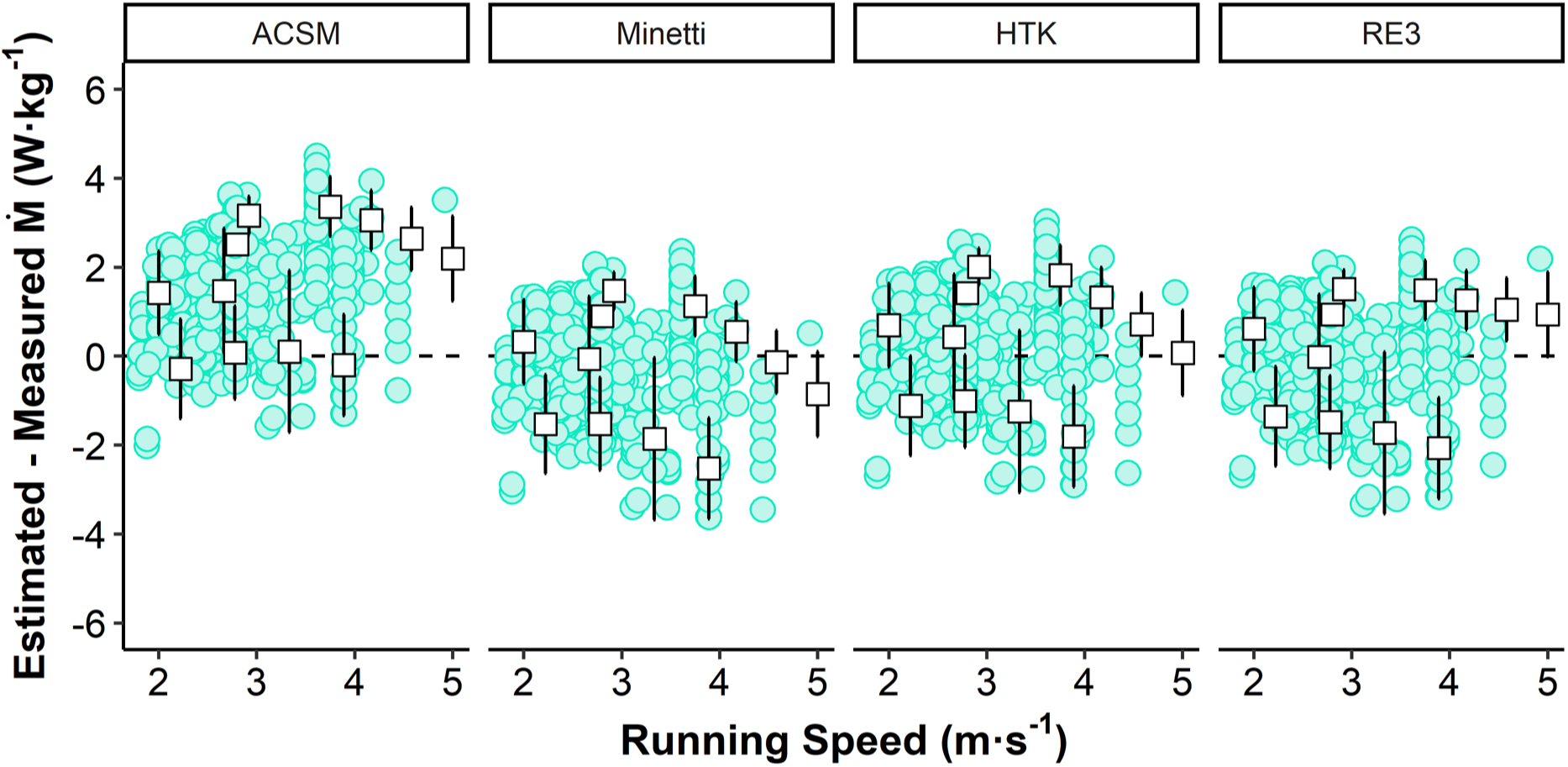
Paired differences between estimated and measured rates of metabolic rate energy expenditure (Ṁ) during level running across running speed for individual subject data (green circles) and group mean data (white squares) ± standard deviation. ACSM, American College of Sports Medicine running equation; Minetti, energy cost of running equation from Minetti et al. (2002); HTK, Hoogkamer-Taboga-Kram equation; RE3, Running Energy Expenditure Estimation graded running equation.

**Fig 3.**
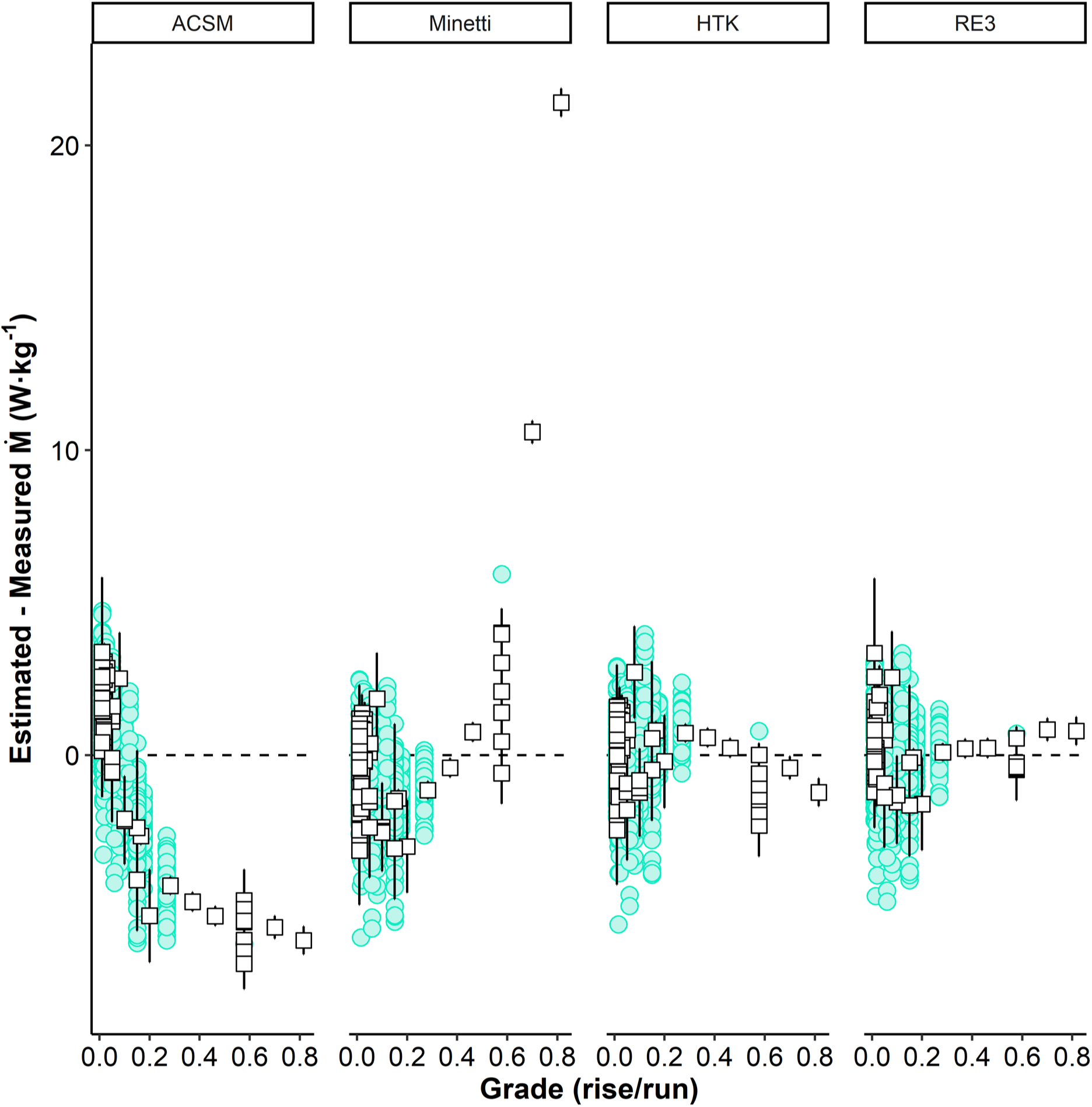
Paired differences between estimated and measured rates of metabolic rate energy expenditure (Ṁ) during uphill running across grade for individual subject data (green circles) and group mean data (white squares) ± standard deviation. ACSM, American College of Sports Medicine running equation; Minetti, energy cost of running equation from Minetti et al. (2002); HTK, Hoogkamer-Taboga-Kram equation; RE3, Running Energy Expenditure Estimation graded running equation.

**Fig 4.**
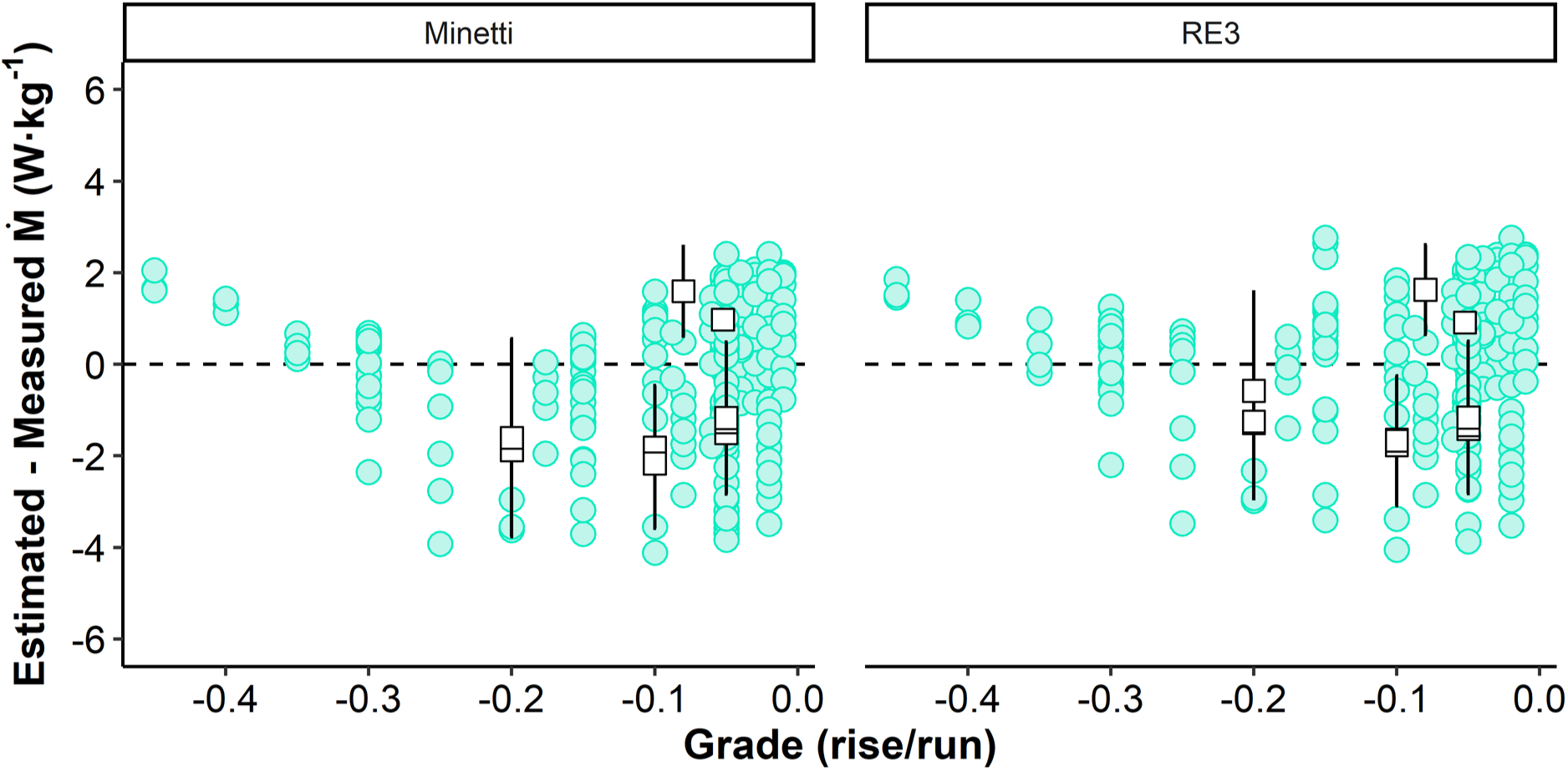
Paired differences between estimated and measured rates of metabolic rate energy expenditure (Ṁ) during downhill running across grade for individual subject data (green circles) and group mean data (white squares) ± standard deviation. Minetti, energy cost of running equation from Minetti et al. (2002); RE3, Running Energy Expenditure Estimation graded running equation.

**Table 3.** Accuracy and precision of graded running equations. ACSM, American College Sports Medicine running equation; Green circles, individual subject data; HTK, Hoogkamer-Taboga-Kram equation; Minetti, energy cost of running equation from Minetti et al. (2002); RE3, Running Energy Expenditure Estimation graded running equation; RMSD, root-mean-square deviation; SD, standard deviation of paired differences; White squares, group mean ± standard deviation.

| Grade | Equation | Difference ($\text{W}\cdot\text{kg}^{-1}$) | | Percent Difference | |
| --- | --- | --- | --- | --- | --- |
| | | Bias ± SD | RMSD | Bias ± SD | RMSD |
| Level | ACSM | $1.35 \pm 1.22$ | 1.82 | $11.6 \pm 10.5$ | 15.6 |
| | Minetti | $-0.53 \pm 1.34$ | 1.44 | $-2.4 \pm 9.4$ | 9.7 |
| | HTK | $-0.08 \pm 1.30$ | 1.30 | $1.0 \pm 9.7$ | 9.7 |
| | RE3 | $-0.26 \pm 1.24$ | 1.27 | $-0.6 \pm 9.2$ | 9.2 |
| Uphill | ACSM | $-0.25 \pm 2.16$ | 2.17 | $-0.4 \pm 13.1$ | 13.1 |
| | Minetti | $-0.67 \pm 2.07$ | 2.18 | $-3.4 \pm 11.7$ | 12.2 |
| | HTK | $0.04 \pm 1.45$ | 1.45 | $0.8 \pm 8.7$ | 8.7 |
| | RE3 | $-0.01 \pm 1.41$ | 1.41 | $0.5 \pm 8.6$ | 8.6 |
| Downhill | Minetti | $-0.39 \pm 1.52$ | 1.57 | $-1.8 \pm 15.0$ | 15.1 |
| | RE3 | $0.02 \pm 1.45$ | 1.45 | $1.4 \pm 14.6$ | 14.7 |

## Discussion

Using a large dataset compiled from 26 studies with individual participant data and 12 studies reporting group mean data, we fit a new graded running term for the RE3 model that enables Ṁ estimation for level, uphill, and downhill running. Using this same large, compiled dataset, we fit an updated formula for the physiologically-based HTK equation for level and uphill running. Both methods proved to be accurate and precise estimators of running energetics on various slopes with demonstrated improvements over two of the most used estimation equations. Consequently, we provide a publicly available web-based metabolic rate calculator that simplifies estimation for researchers, practitioners, and runners alike (University of Massachusetts Integrative Locomotion Lab., 2025).

As can be seen in Fig3, on steeper inclines, the ACSM equation underestimates Ṁ whereas the Minetti equation overestimates Ṁ. Perhaps this point can be best illustrated by considering an athlete running in a vertical kilometer (VK) race. The official VK record for is 28 min 53 s set in Fully, Switzerland (Vertical Kilometer World Circuit., 2025). That course ascends 1,000 m over a distance of 1,920 m equating to an average grade of 0.52 and an average running speed of 1.11 m·s^-1^. The ACSM equation predicts Ṁ to be 16.99 W·kg^-1^ whereas the Minetti equation predicts 27.92 W·kg^-1^, a nearly two-fold range. Whereas the RE3 model (25.51 W·kg^-1^) and new HTK model (24.13 W·kg^-1^) predict similar intermediate Ṁ values.

The novel graded running term developed for the RE3 model allows for estimating Ṁ across an extreme range of downhill and uphill grades (−0.45 to 0.82), the largest range of surface grades that any equation has been developed on to date. This is useful not only for applications such as mountain running (Balducci et al., 2016; Vernillo et al., 2020) but also for team field sports where the energy cost of accelerated running has been estimated based on the energy cost of running of an “equivalent slope”, or a slope with a comparable angle of forward body lean (Osgnach et al., 2010). For the RE3 model to be most applicable to military populations, there is a need to characterize the metabolic effects of the loads added by body armor or utility vests (Looney et al., 2024), backpacks/rucksacks (Looney et al., 2022), military footwear such as combat boots (Lavoie et al., 2023), and other operationally-relevant equipment (Soucy et al., 2023) that servicemembers may be required to carry while running or sprinting.

Hoogkamer et al. (2014) expanded the cost of generating force hypothesis for level running to uphill running and observed efficiency values for producing work to vertically lift the center of mass only slightly higher than physiologically realistic values. However, the accuracy of the original HTK model beyond the speeds (up to 3.0 m·s^-1^) and the inclines (up to 8°, 0.14 grade) measured to fit the equation had not been assessed. By fitting the HTK equation to a much larger data set including faster running speeds and steeper grades, we were able to update the equation within the original biomechanical framework. The best fit resulted in an efficiency value for producing work to vertically lift the center of mass of 23.0%, which is physiologically realistic, more so than the 29.4% of the original equation. Intriguingly, the accuracy and precision of this updated equation is similar to that of the phenomenological RE3 equation. As such the updated HTK model provides a framework linking running biomechanics and energetics with many opportunities. It allows for personalized Ṁ estimates based on individual measures of standing Ṁ and efficiency, and it can be further expanded to include downhill running and load carriage, all based on biomechanical and muscle physiological theory.

As they are designed for simple, generalized applications, the HTK and RE3 equations estimate Ṁ for an average individual and do not account for all the variability in mass-specific running economy across the spectrum of runners (Daniels, 1985). Thus, erring on the side of overconsuming calories for competitive, recreational, or tactical athletes during an event would be prudent. While running Ṁ can be individualized based on key aerobic fitness parameters, there are many to choose from and most can only be measured accurately using laboratory equipment that can be difficult for many runners to access. Additionally, these aerobic fitness parameters do not perfectly correlate with inter-individual variability in Ṁ as even trained runners with similar maximal oxygen uptake (V̇O_2max_) have been shown to vary in running economy by upwards of 30% (Barnes & Kilding, 2015; Daniels, 1985). Furthermore, running economy does not always translate between different conditions, such as running on the level versus steep slopes (Lemire et al., 2021), highlighting the importance of exercise familiarity and training experience. Developing an easy-to-use method for calculating individualized Ṁ estimates that can accommodate a variety of aerobic fitness parameters as well as account for experience and familiarity with specific types of running is a difficult but important aim for future research.

While Vernillo et al. (2017) provide a concise summary of what is known about the biomechanics and physiology of downhill running, developing a mechanistic-based equation for downhill running will require a deeper understanding especially for grades steeper than 15% (−9°). Even just quantifying the relevant metabolic rate for downhill running is not straightforward because V̇O_2_ (and presumably Ṁ) gradually increases by as much as 10% over trial durations of 40-45 min down −10% grades (Dick & Cavanagh, 1987; Westerlind et al., 1994). Westerlind et al. (1994) found that in a second downhill running trial, initial V̇O_2_ was ∼ 8.5% lower than the initial value in the first trial but then increased again (by ∼ 8%) over the course of the second 40 min trial. Note that Dick and Cavanagh (1987) studied experienced runners whereas Westerlind et al. (1994) intentionally studied individuals who were not accustomed to downhill running. Future research is needed to determine how leg muscle fascicles actually behave during downhill running beyond the usual assumption of eccentric action. Given that most road-racing courses have moderate inclines and declines, future studies should quantify the metabolic cost of downhill running at grades < 10% with finer resolution so that the shape of the grade vs. Ṁ relationship can be better characterized. A major hindrance to downhill running research is the scarcity of suitable commercial treadmills. Treadmill modifications are necessary to maintain belt speed post-foot contact, and potential motor damage is a concern. Adding an inertial flywheel can certainly help (Kram et al., 1998). Smart treadmill motor controllers that surge power at foot strike is another alternative.

## Conclusion

Our analysis provides researchers, practitioners, and runners with the capability to estimate Ṁ during running on level, uphill, and downhill slopes using the novel graded running term fit for the existing RE3 model. The newly developed web browser-based metabolic rate calculator (University of Massachusetts Integrative Locomotion Lab., 2025) provides users with a simple way to calculate Ṁ with the updated RE3 model. Additionally, the enhanced, generalized version of the HTK equation provides close estimates of Ṁ during level and uphill running that is consistent with present understanding of level and uphill running biomechanics and muscle physiology.

**Appendix 1** Deriving the efficiency value assumed by the American College of Sports Medicine (ACSM) running equation.

The ACSM equation for estimating oxygen uptake (V̇O_2_) during running is:

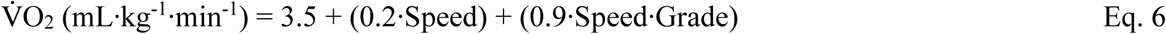

where speed is in units of m/min and grade is expressed as a decimal percentage, i.e. 10% grade is 0.10. The vertical mechanical power (work rate) in watts (W) is equal to the product of body mass, gravitational acceleration (9.81 m/s^2^), treadmill belt velocity (in m/s) and the sin of the incline angle (θ). For example, the vertical mechanical power for a 70 kg person running at 180 m/min up a 10% incline (θ = 5.7°) is equal to mg v sin θ or 70 kg·9.81 m·s^-2^·3 m·s^-1^·sin 5.7° which is equal to 204 W. The corresponding V̇O_2_ from the third term in the ACSM equation for is 0.9·180·0.10 or 16.2 mL O_2_/kg/min. Multiplying 16.2 by body mass (70 kg) and dividing by 60 s per min yields 18.9 mL O_2_·s^-1^ which converts to a rate of metabolic energy expenditure (Ṁ) of 395 W, assuming 20.9 J/mL O_2_ (Gill et al. 2023). Efficiency is the ratio of mechanical power and Ṁ which, in this example, is 204 W/395 W or 51.9%.

## Acknowledgments

We thank Debanshi Jain for her support in developing the website-based metabolic rate estimation application.

## Abbreviations

ACSM: American College of Sports Medicine
COM: Center of mass
CoT: Metabolic cost of transport
Eff_vCOM_: Efficiency of producing mechanical work to lift the center of mass against gravity
G: Surface grade
GRF: Ground reaction force
HTK: Hoogkamer-Taboga-Kram
iOS: iPhone Operating System
Ṁ: Rate of metabolic energy expenditure
RE3: Running Energy Expenditure Estimation
RER: Respiratory exchange ratio
RMSD: Root-mean-square deviation
S: Running speed
SD: Standard deviation
t_c_: Time of foot-ground contact
V̇CO_2_: Rate of carbon dioxide production
V̇O_2_: Rate of oxygen uptake

## Statements and Declarations

No funding was received for conducting this study.

